# Visual response characteristics in lateral and medial subdivisions of the rat pulvinar

**DOI:** 10.1101/2020.01.26.920454

**Authors:** Andrzej T. Foik, Leo R. Scholl, Georgina A. Lean, David C. Lyon

## Abstract

The pulvinar is a higher-order thalamic relay and a central component of the extrageniculate visual pathway, with input from the superior colliculus and visual cortex and output to all of visual cortex. Rodent pulvinar, more commonly called the lateral posterior nucleus (LP), consists of three highly-conserved subdivisions, and offers the advantage of simplicity in its study compared to more subdivided primate pulvinar. Little is known about receptive field properties of LP, let alone whether functional differences exist between different LP subdivisions, making it difficult to understand what visual information is relayed and what kinds of computations the pulvinar might support. Here, we characterized single-cell response properties in two V1 recipient subdivisions of rat pulvinar, the rostromedial (LPrm) and lateral (LPl), and found that a fourth of the cells were selective for orientation, compared to half in V1, and that LP tuning widths were significantly broader. Response latencies were also significantly longer and preferred size more than three times larger on average than in V1; the latter suggesting pulvinar as a source of spatial context to V1. Between subdivisons, LPl cells preferred higher temporal frequencies, whereas LPrm showed a greater degree of direction selectivity and pattern motion detection. Taken together with known differences in connectivity patterns, these results suggest two separate visual feature processing channels in the pulvinar, one in LPl related to higher speed processing which likely derives from superior colliculus input, and the other in LPrm for motion processing derived through input from visual cortex.

**Significance Statement:** The pulvinar has a perplexing role in visual cognition as no clear link has been found between the functional properties of its neurons and behavioral deficits that arise when it is damaged. The pulvinar, called the lateral posterior nucleus (LP) in rats, is a higher order thalamic relay with input from the superior colliculus and visual cortex and output to all of visual cortex. By characterizing single-cell response properties in anatomically distinct subdivisions we found two separate visual feature processing channels in the pulvinar, one in lateral LP related to higher speed processing which likely derives from superior colliculus input, and the other in rostromedial LP for motion processing derived through input from visual cortex.

## Introduction

In the mammalian visual system, there are two major pathways for visual information to reach visual cortex. The geniculostriate pathway carries retinal signals through the lateral geniculate nucleus (LGN) to primary visual cortex (V1), and accounts for the majority of the visual information flow (Jones, 1985). The extrageniculate pathway relays retinal input through the superior colliculus then pulvinar to all of visual cortex; it’s function is less well understood (Waleszczyk et al., 1999, 2004; Lyon et al., 2010). This is especially true for rodent models where little has been reported on receptive field properties of mouse pulvinar (a.k.a. lateral posterior nucleus; LP; Roth et al., 2015; Ahmadlou et al., 2018; Bennett et al., 2019), and even less so in rats (Montero et al., 1968).

Anatomical similarities across species suggest a general organization of pulvinar unchanged over evolutionary time scales ((Lyon et al., 2003; Zhou et al., 2017); see Figure 1). Rodent pulvinar consists of three highly-conserved subdivisions, and offers the advantage of simplicity in its study compared to the more subdivided primate pulvinar. In rat and mouse, the pulvinar is divided based on cytoarchitecture and connectivity into caudomedial (LPcm), rostromedial (LPrm), and lateral (LPl) (see Figure 1AB; (Takahashi, 1985; Nakamura et al., 2015)). These subdivisions are homologous to the three found in squirrel (Figure 1D), a highly visual rodent (Robson and Hall, 1977; Baldwin et al., 2011), and tree shrew (Figure 1C), a close primate relative (Lyon et al., 2003); and share anatomical similarities to dorsomedial (PLdm), ventrolateral (PLvl), and inferior (PI) subdivisions of primate pulvinar (Figure 1F; (Lyon et al., 2003; Kaas and Lyon, 2007).

**Figure 1.**
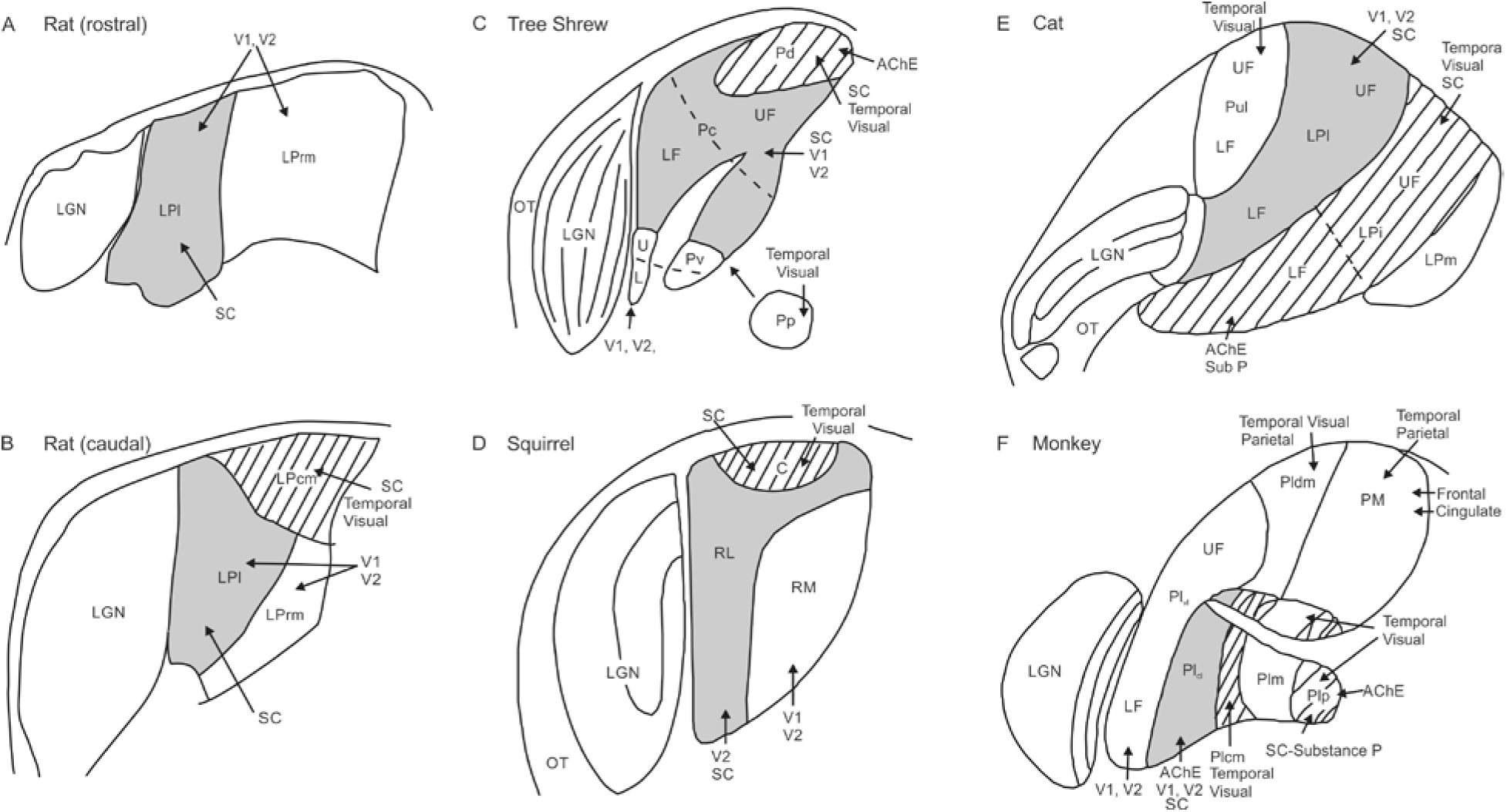
Schematic of pulvinar anatomy across five species. Regions receiving dense superior colliculus (SC) input and with connections to temporal visual cortex (hatched) include caudomedial pulvinar (LPcm) in rat (B), dorsal pulvinar (Pd) in tree shrew (C), caudal pulvinar (C) in squirrel (D), intermediate LP (LPi) in cat (E), and inferior pulvinar (PI) in monkey (F), although not all of PI receives input from SC. Regions receiving sparse input from superior colliculus with connections to V1 and V2 (shaded grey) include lateral pulvinar (LPl) in rat (A, B), caudal pulvinar in the tree shrew (Pc; C), rostrolateral pulvinar (RL) in squirrel (D), lateral LP in cat (Lpl; E), and caudolateral inferior pulvinar (PIcl) in monkey (F). Regions with visual cortex connections only (no SC input) include rostromedial pulvinar (LPrm) in rat (A,B), ventral pulvinar (Pv) in tree shrew (C), rostromedial pulvinar (RM) in squirrel (D), lateral LP (LPl) in cat (E), and ventrolateral lateral pulvinar (PLvl) in monkey (F). Additional subregions include posterior pulvinar (Pp) in tree shrew, the pulvinar (Pul) and medial LP (LPm) in cat, and the medial (PM) and dorsomedial lateral (PLdm) pulvinar in monkey. Optic tract (OT) and lateral geniculate nucleus (LGN) are shown for reference. Adapted from Lyon et al. (2003) and Nakamura (2015); not to scale.

The pulvinar receives significant corticothalamic modulatory efferents (Benevento and Rezak, 1976), and sends many modulatory projections to all of visual cortex (Trageser and Keller, 2004), making it well suited to mediate information transfer between cortical areas. Recent behavioral studies in monkey have implicated it in directing visually guided actions (Wilke et al., 2010), and projections from pulvinar to V1 in mice have been shown to carry information about discrepancy between visual and motor inputs (Roth et al., 2016). Further work in mouse has demonstrated that pulvino-recipient neurons in higher cortical visual areas respond strongly to pulvinar activity if they in turn project to areas involved in guiding visual movement (Zhou et al., 2018).

Single unit recordings in primate pulvinar identified some basic receptive field properties that highlighted important differences between some subdivisions. For example, cells in the dorsomedial pulvinar (Pdm) have much larger receptive fields, longer response latencies, and are less retinotopically organized than cells in inferior (PI) and lateral (PL) pulvinar (Petersen et al., 1985); cells in the latter are also more selective for orientation and direction (Gattass et al., 1979; Bender, 1982; Petersen et al., 1985). These differences could contribute to the respective role of each subdivision in guiding visual behavior, since receptive fields with faster response latencies are more suited for motion processing and dorsal stream vision whereas cells with higher spatial acuity are more likely to process ventral stream object vision (Shetht and Young, 2016).

In mouse or rat, little is known about receptive field properties, let alone whether functional differences exist the different LP subdivisions (Montero et al., 1968; Roth et al., 2015; Ahmadlou et al., 2018; Bennett et al., 2019). This makes it difficult to predict what role pulvinar regions have in modulating cortical activity and whether rodent LP would be a useful model for primate pulvinar. In order to address this lack of single unit data, we collected and compared spatiotemporal receptive fields of neurons in LPrm and LPl subdivisions in the rat and compared them with V1 responses. While both regions receive significant input from visual cortex, LPl also receives superior colliculus input. We hypothesized that different connectivity between the two subdivisions would be reflected in different profiles of visual information processing.

## Materials and Methods

### Animals

The experiments were performed on 21 adult Long-Evans rats of either sex. Animals were treated in accordance with the NIH guidelines for the care and use of laboratory animals, the ARVO Statement for the Use of Animals in Ophthalmic and Vision Research, and under a protocol approved by the Institutional Animal Care and Use Committee of UC Irvine.

### Experimental procedures

Rats were initially anesthetized with 2% isoflurane in a mixture of N_2_O/O_2_ (70%/30%), then placed into a stereotaxic apparatus. Animals were given dexamethasone (0.4 ml/kg, Dexamethasone) injection prior to surgery and painkiller injection after surgery (0.05 ml/kg, FluMeglumine). A small, custom-made plastic chamber was glued to the exposed skull. After one day of recovery, re-anesthetized animals were placed in a custom-made hammock, maintained under isoflurane anesthesia (2% in a mixture of N_2_O/O_2_), and a single tungsten electrode was inserted into a small craniotomy above visual cortex. Once the electrode was inserted, the chamber was filled with sterile saline and sealed with sterile wax. During recording sessions, animals were kept sedated under light isoflurane anesthesia (0.2 – 0.4%) in a mixture of N_2_O/O_2_. EEG and EKG were monitored throughout the experiments and body temperature was maintained with a heating pad (Harvard Apparatus, Holliston, MA, USA). Following several recording sessions, rats were deeply anesthetized with isoflurane and sodium pentobarbital, then perfused transcardially first with saline, then 4% paraformaldehyde. Brains were removed and stored in 30% sucrose in phosphate buffer for histology.

### Single unit recordings, visual stimulation and data acquisition

Single unit recordings were made with one or multiple perpendicularly inserted single tungsten electrodes. Electrodes were coated with 1,1′-dioctadecyl-3,3,3′,3′-tetramethyl-indocarbocyanine perchlorate fluorescent dye (DiI, Sigma; (DiCarlo et al., 1996)). Multichannel recordings were acquired using a 32-channel Scout recording system (Ripple, Salt Lake City, UT, USA). Signals containing spikes were bandpass filtered from 500 Hz to 7 kHz and stored on a computer hard drive at 30 kHz sampling frequency. Spikes were sorted online in Trellis software (Ripple, Salt Lake City, UT, USA) while performing visual stimulation. Visual stimuli were generated in Matlab (Mathworks, USA) using Psychophysics Toolbox (Brainard, 1997; Pelli, 1997; Kleiner et al., 2007) and displayed on a gamma-corrected LCD monitor (55 inches, 60 Hz; RCA, NY, USA) at 1920×1080 pixels resolution and 52 cd/m^2^ mean luminance. Stimulus onset times were corrected for LCD monitor delay using a photodiode and microcontroller (in-house design based on Arduino microcontroller). Visually responsive cells were found using either 100% contrast drifting grating stimuli or brief (500 ms) flashes of white on a black screen. Responsive cells were tested with drifting gratings in order to determine receptive field location and optimal parameters for orientation/direction, spatial/temporal frequency, aperture, contrast, etc.

### Histology

Brains were sectioned coronally at 40 μm on a freezing microtome, stained for DAPI, and observed under fluorescent and bright-field light using a Zeiss Axioplan2 microscope (White Plains, NY). Digital images of electrode tracks stained with DiI fluorescent dye (Figure 2A) were captured using a low-light-sensitive video camera (Cooke Sensicam QE) and appropriate filters using Neurolucida software (MBF Bioscience, Williston, VT USA). Brain sections were visually inspected for the location of DiI fluorescent electrode tracts. Tract penetrations were determined to be in either the LPl or LPrm subdivision based on myeloarchitecture seen in autofluorescence and DAPI staining, relative position to other tracts, and stereotaxic coordinates according to the rat brain atlas by Paxinos and Watson (2013; see Figure 2B-D).

**Figure 2.**
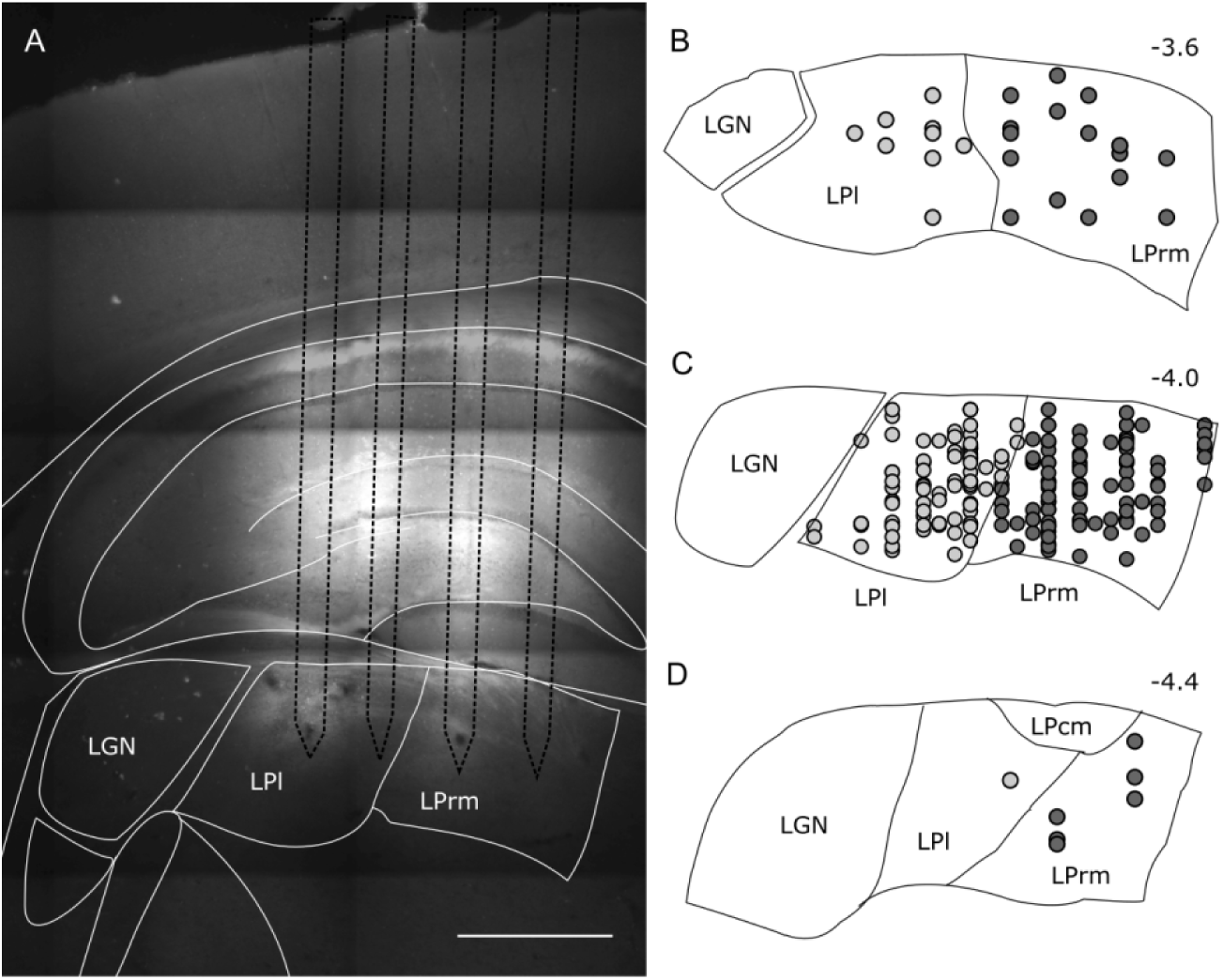
Location of recordings sites in lateral (LPl) and rostromedial (LPrm) subdivisions of the pulvinar. (A) Brain section showing four electrode marks labeled with DiA targeting LPl and LPrm. (B-D) Most of the recording sites (filled circles) were localized between −3.6 and −4.4 mm from Bregma, and centered around −1.5 mm laterally from the midline for LPrm (dark grey circles), and −2.5 mm laterally for LPl (light grey circles).

### Data analysis and statistics

Tuning curves were constructed using the mean firing rate during stimulus presentation, averaged over multiple repetitions (8 for grating stimuli, 100 – 300 for flash stimulus). Optimal parameters were determined based on the maximum firing rate.

Responses were considered as selective if preference to any stimulus condition was present in the tuning curve for given stimulus parameter and response was higher than background activity. The percentages in Table 1 were calculated according to the total number of cells recorded for each subdivision.

**Table 1.** Percentage of visually responsive and selective neurons in LP subdivisions and V1.

|  | ***Total***<br>***Number*** | ***Visual*** | ***Grating*** | ***Orientation*** | ***Size*** |
| --- | --- | --- | --- | --- | --- |
| ***LPrm*** | 241 | 79 | 52 | 26 | 28 |
| ***LPI*** | 229 | 74 | 43 | 19 | 17 |
| ***V1*** | 101 | 88 | 79 | 54 | 62 |

Peri-stimulus time histograms (PSTH) were calculated with 10 ms bin width. Response latency was defined by the time at which the instantaneous firing rate exceeded mean background activity calculated from 100 ms before stimulus onset plus two standard deviations of each PSTH, and corrected manually.

Orientation selectivity index (OSI) was calculated as follows:

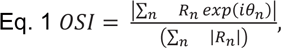

where θ*_n_* is the *n*th orientation of the stimulus and R_n_ is the corresponding response.

Direction selectivity (DSI) index was calculated according to Eq. 1 but where θ*_n_* is the *n*th direction of the grating drift and R_n_ is the corresponding response.

Orientation tuning bandwidth was calculated by fitting orientation responses to double Gaussian distributions (Carandini and Ferster, 2000; Alitto and Usrey, 2004) using:

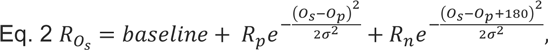

where O_s_ is the stimulus orientation, R_Os_ is the response to different orientations, O_p_ is the preferred orientation, R_p_ and R_n_ are the responses at the preferred and non-preferred direction, σ is the tuning width, and ‘baseline’ is the offset of the Gaussian distribution. Gaussian fits were estimated without subtracting spontaneous activity, similar to the procedures of Alitto & Usrey (Alitto and Usrey, 2004). The orientation tuning bandwidth of each tuning curve was measured in degrees as the half-width at half-height (HWHH), which equals 1.18 × σ based on the equation above.

Size tuning curves were fitted by a difference of Gaussian (DoG) function:

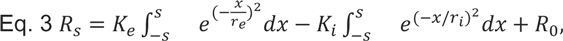

in which R_s_ is the response evoked by different aperture sizes. The free parameters, K_e_ and re, describe the strength and the size of the excitatory space, respectively; Ki and ri represent the strength and the size of the inhibitory space, respectively; and R_0_ is the spontaneous activity of the cell.

The optimal spatial and temporal frequency was extracted from the data fitted to Gaussian distribution using the following equation (DeAngelis et al., 1995; Van Den Bergh et al., 2010):

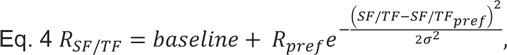

Where R_SF/TF_ is estimated response, R_pref_ indicates response at a preferred spatial or temporal frequency, SF/TF indicates spatial or temporal frequency, σ is the standard deviation of the Gaussian and baseline is Gaussian offset.

Tuning curve fits and goodness of fits were calculated using the EzyFit curve fitting toolbox (http://www.fast.u-psud.fr/ezyfit/).

### Classification of pattern motion and component motion selective cells

The responses of LP cells were classified according to classical approach using partial correlations formula (Movshon et al., 1985; Casanova and Savard, 1996; Movshon and Newsome, 1996; Merabet et al., 1998; Palagina et al., 2017):

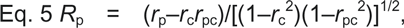

where *R*_p_ represents the partial correlation coefficient for the pattern prediction, *r*_c_ is the correlation coefficient of the grating motion response and the component motion (CM) prediction, *r*_p_ is the correlation coefficient for the grating response and pattern motion (PM) response, and *r*_pc_ is the correlation coefficient for the pattern motion tuning curve and CM prediction. The *R*_c_ is the partial correlation defined for the CM prediction and is calculated by exchanging *r*_p_ with *r*_c_ in the equation. A cell is considered pattern motion-selective when the value of *R*_p_ is significantly greater than either *R*_c_ or zero. The CM prediction is calculated as the average of the grating direction tuning curve shifted by the offset of pattern motion components (in this case it is 90°).

Differences were considered significant at p ≤ 0.05 for the two-tailed Mann-Whitney U-test for comparison between groups, Wilcoxon signed-rank test for comparison within each group and Kolmogorov-Smirnov (K-S test) for distribution differences. Error bars indicate the standard error of the mean (SEM). All offline data analysis and statistics were performed using Matlab (Mathworks, USA).

## Results

To determine response characteristics in two rostral subdivisions of the pulvinar, or, lateral posterior thalamic nucleus (LP), we recorded from 229 and 241 neurons in LPl and LPrm, respectively, and then compared these to responses of 101 V1 neurons in 21 Long-Evans pigmented rats. A summary of reconstructed LP recording sites is shown in Figure 2B-D. Recordings in LPl were confined to be around −4 mm from Bregma (M = −3.9 ± 0.1 mm) and 2.5 mm from the midline (M = 2.6 ± 0.3 mm) and in LPrm to be around −4 mm from Bregma (M = −4.0 ± 0.2 mm) and 1.5 mm from the midline (M = 1.8 ± 0.3 mm).

In V1, most cells were visually responsive and easy to find using high contrast grating stimuli (Table 1). In LP, cells had less robust responses to gratings but were readily identified using flash stimuli. Significantly fewer cells were visually responsive in both LPrm (*χ*^2^(1) = 4.1, P < 0.05) and LPl (*χ*^2^(1) = 8.4, P < 0.01) compared to V1 (Table 1). Visually evoked responses of neurons to brief (500 ms) flashes of light also revealed differences in latency between LP and V1 neurons (Figure 3 A-G). Both LPrm (P< 0.001) and LPl (P = 0.001) had significantly longer latencies than V1 on average (Figure 3G), although the shortest response latencies were observed in LP rather than V1 (Figure 3AC). Mean spontaneous firing rate was also significantly higher in LPrm (P < 0.02) and LPl (P < 0.01) compared to V1 (Figure 3 BDFG).

**Figure 3.**
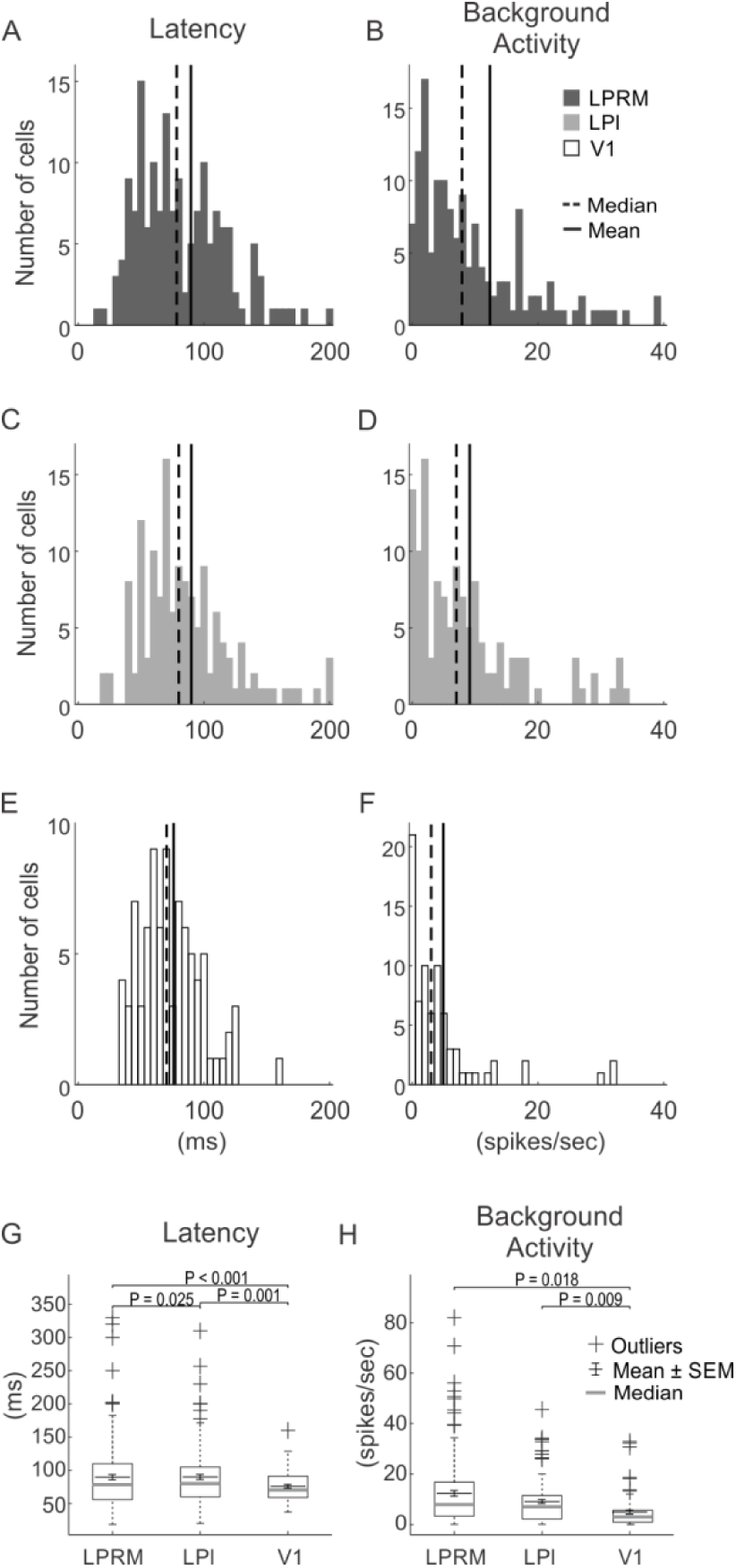
Population summary for measures of response latency and background activity in pulvinar and V1. The distribution of response latency (A,C,E) and spontaneous background activity (B,D,F) of cells recorded from LPrm (A,B), LPl (C,D), and V1 (E,F); vertical solid and dashed line show mean and median for each distribution, respectively. Whisker plots show comparisons of mean and median responses for latency (G) and background activity (H).

### Spatial and temporal properties of single neurons in LPrm, LPl, and V1

All visually responsive cells were tested for optimal sinusoidal grating parameters such as direction, orientation, spatial frequency, size, temporal frequency, and contrast. In LP, about one third of the cells responsive to flash stimuli did not have strong responses to drifting gratings (Table 1), even with low spatial frequencies (0.001 °/cycle), and were excluded from further analysis. In V1, about 10% of the cells were also excluded for not responding to grating stimuli. Representative examples of grating responsive neurons are shown in Figure 4. For these cells, spatial frequency responses for LPrm (Figure 4G), LPl (Figure 4H), and V1 (Figure 4I) showed similar tuning profiles with preferences well over 0.01 cycle/°. Likewise, in the population, average optimal spatial frequencies were also very similar for all tested regions measuring between 0.05 and 0.06 cycle/° (Figure 5A, D, G, J).

**Figure 4.**
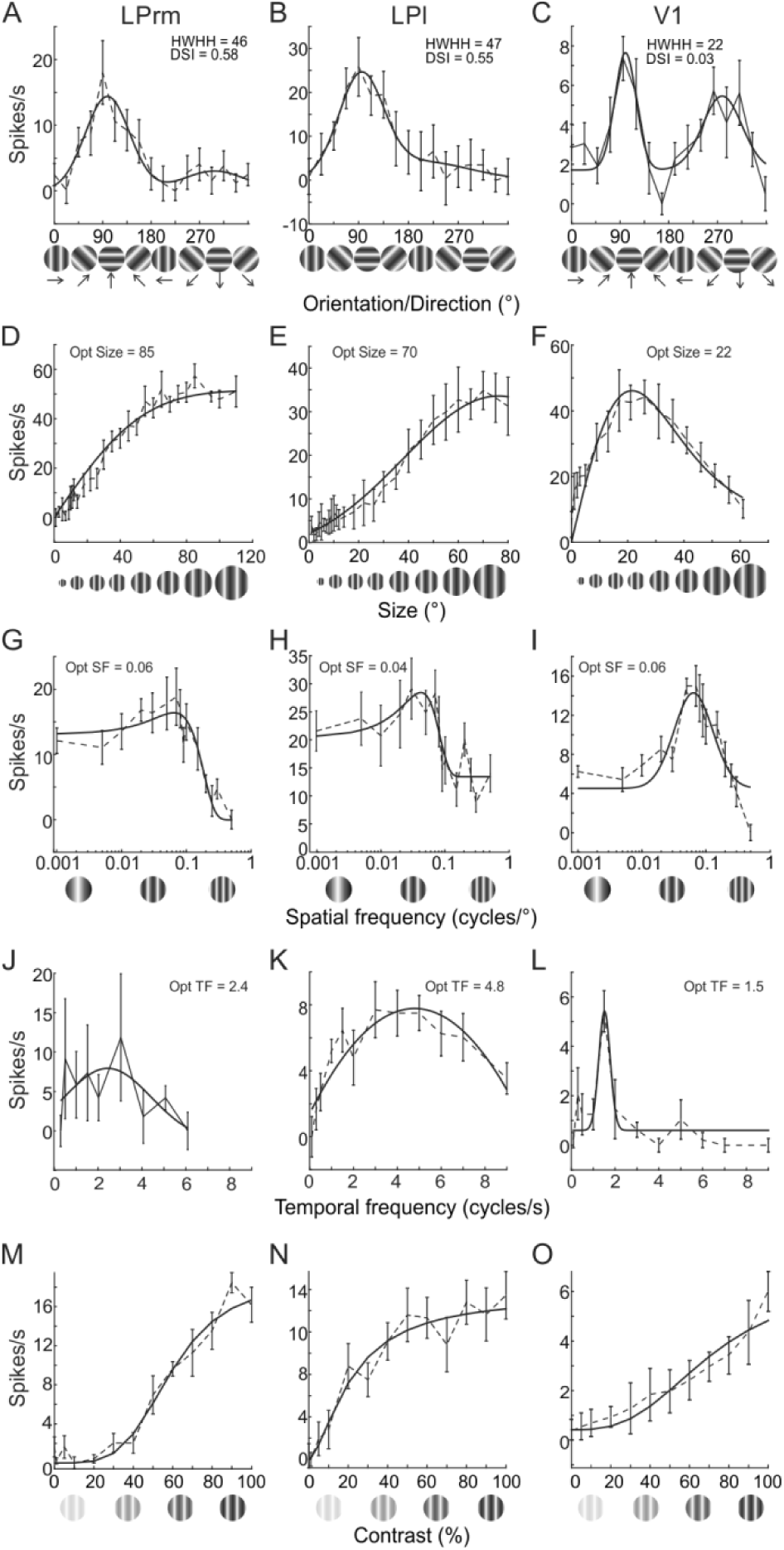
Representative examples of spatiotemporal response properties of pulvinar and V1 cells. One cell each in LPrm (left column), LPl (middle column) and V1 (right column), before (dotted lines) and after (solid lines) curve fitting. In (A-C), orientation tuning bandwidth was calculated based on the half width at half height (HWHH) of the preferred direction, and direction selectivity index (DSI) was also computed. Size (D-F), spatial frequency (G-I), temporal frequency (J-L) were calculated based on the value realizing the highest firing rate. Contrast response curves (M-O) were also generated.

**Figure 5.**
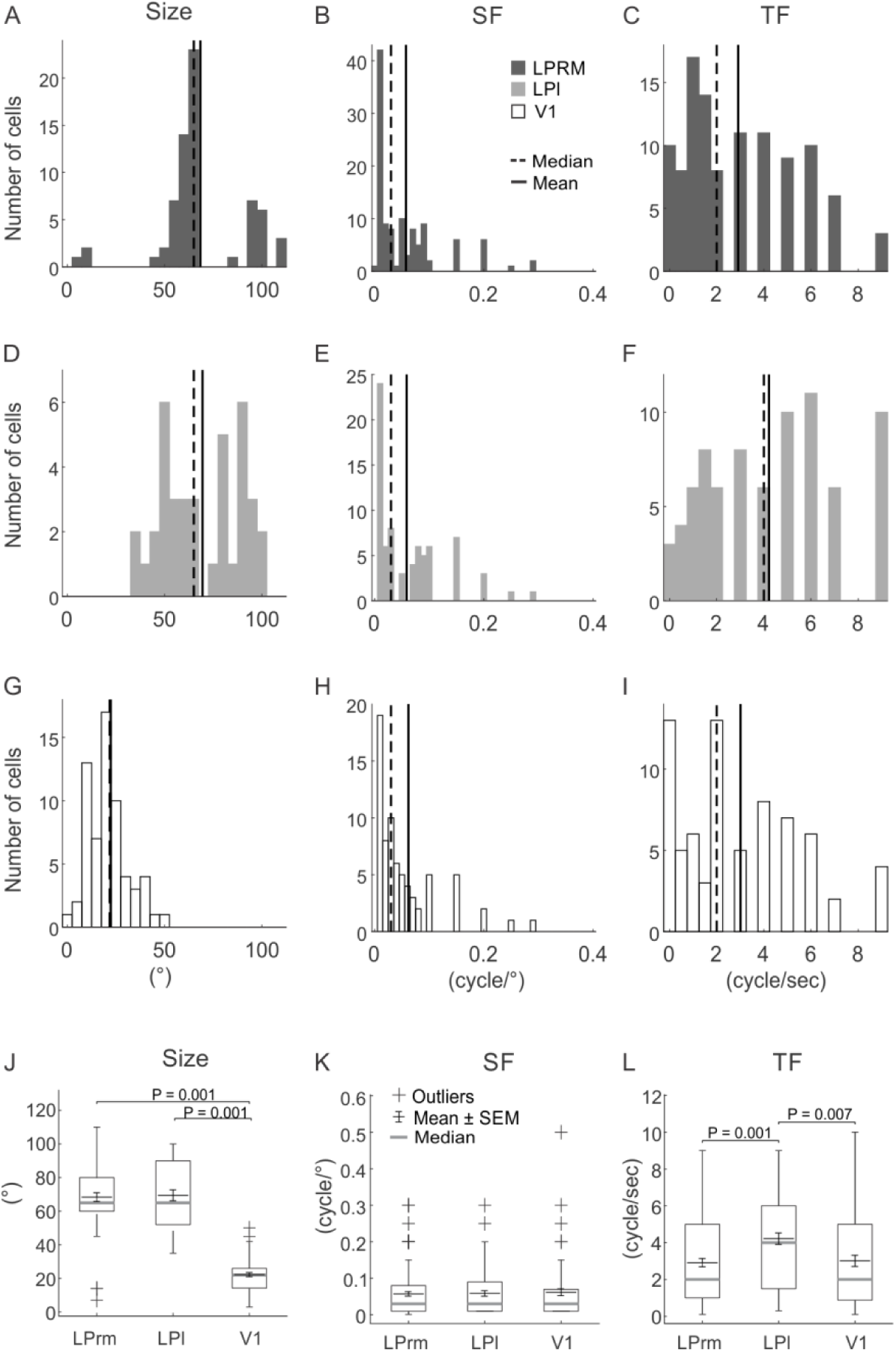
Population summary of optimal parameters for pulvinar and V1 cells. The distributions of optimal size (A-G), spatial frequency (SF; B-H) and temporal frequency (TF; C,F,I) are shown for LPrm (A-C), LPl (D-F) and V1 (G-I). Whisker plots provide comparisons of mean and median responses of optimal size (J), spatial (K), and temporal (L) frequencies.

Orientation tuning, on the other hand, was generally broader in LP compared to V1. For example, tuning widths (HWHH) were similar for the LPrm (46°; Figure 4A) and LPl (47°; Figure 4B) cells, and more than double that of the V1 cell (22°; Figure 4C). In contrast, the two LP cells were both highly direction selective (DSI = 0.58 and 0.55) compared to the V1 neuron (DSI = 0.02). Comparing population averages across the three regions, tuning widths were similar between LPrm (47°; Figure 6AJ) and LPl (49°; Figure 6DJ), and significantly broader than V1 (36°, P < 0.01; Figure 5GJ). Likewise, the direction selectivity indices for LPrm (DSI = 0.25; Figure 6BK) and LPl (DSI = 0.23; Figure 6EK) neurons were higher than in V1 (DSI = 0.18; Figure 6HK); however, only the LPrm difference was significant (P < 0.01; Figure 6K). When plotted against the direction selectivity index (DSI) there was a significant correlation with HWHH for LPrm (R = 0.29, P < 0.05; Figure 6C), whereas this correlation was weaker and not significant for LPl (R = 0.1, P > 0.50; Figure 6F) and V1 neurons (R = 0.19, P > 0.15; Figure 6I), indicating a greater degree of direction selectivity in LPrm.

**Figure 6.**
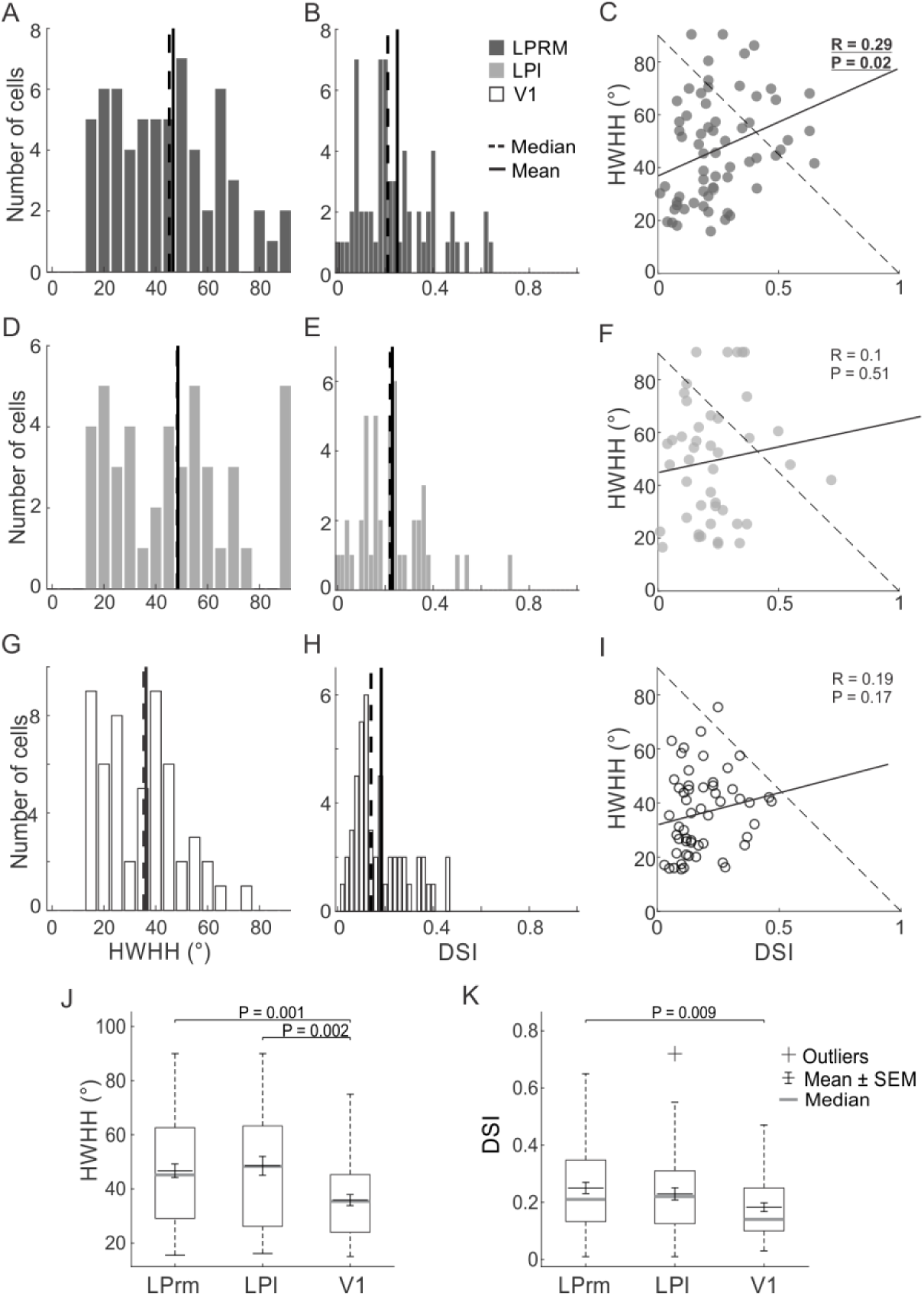
Population summary for measures of orientation and direction of pulvinar and V1 cells. The distributions of HWHH (A,D,G) and DSI (B,E,H) are shown for LPrm (A,B), LPl (D,E) and V1 (GH). Distribution comparisons between HWHH and DSI are shown for LPrm (C), LPl (F) and V1 (I). Whisker plots provide comparisons of mean and median responses for HWHH (J) and DSI (K).

Similar to orientation, both LP subdivisions showed similar size preferences that were substantially larger than for V1. Whereas, the example V1 cell responded strongest to a 22° aperture (Figure 4F), and was subsequently suppressed by larger stimuli, the LPrm (85°; Figure 4D) and LPl (70°; Figure 4E) cells responded best to apertures more than three times larger and plateaued beyond their peak response. These numbers are in agreement with the averages found in the population, where both LPrm (Figure 5A) and LPl (Figure 5D) show more than three times the preferred size (∼70°) compared to V1 (22°, P < 0.001; Figure 5GJ).

The main difference between the two LP subdivisions was found in optimal temporal frequency. In representative examples, the LPl cell had the highest preferred temporal frequency at 4.8 cycle/sec (Figure 4K) compared to the LPrm (2.4 cycle/sec; Figure 4J) and V1 (1.5 cycle/sec; Figure 4L) cells. Similarly, the population average for LPl (4.2 cycle/sec; Figure 5F) was significantly higher than the average temporal frequency of LPrm (2.9 cycle/sec, P = 0.001; Figure 5CL) and V1 (3.0 cycle/sec, P < 0.01; Figure 5IL). Conversely, there was a high similarity in temporal frequency preference between LPrm and V1 neurons (P = 0.96; Figure 5L).

### Neurons in rat LP respond to pattern motion

In addition to component motion of a single drifting grating we also tested a subset of 23 neurons each in LPrm and LPl for detection of pattern motion produced by two drifting gratings set 90° apart. Examples of typical pattern motion tuning curves are presented in Figure 8 for LPrm (top row) and LPl (bottom row). All tested cells had similar response magnitudes to plaid stimuli compared to single gratings (see Figure 8DI). To distinguish pattern motion selectivity, we first used the original pattern vs. component cell classification scheme which measures the correlation between the two (Gizzi et al., 1990; Movshon and Newsome, 1996; Merabet et al., 1998; Ouellette et al., 2004; Smith et al., 2005). This analysis revealed only a few pattern motion selective cells in LPrm (Figure 8A) and none in LPl (Figure 8F).

**Figure 7.**
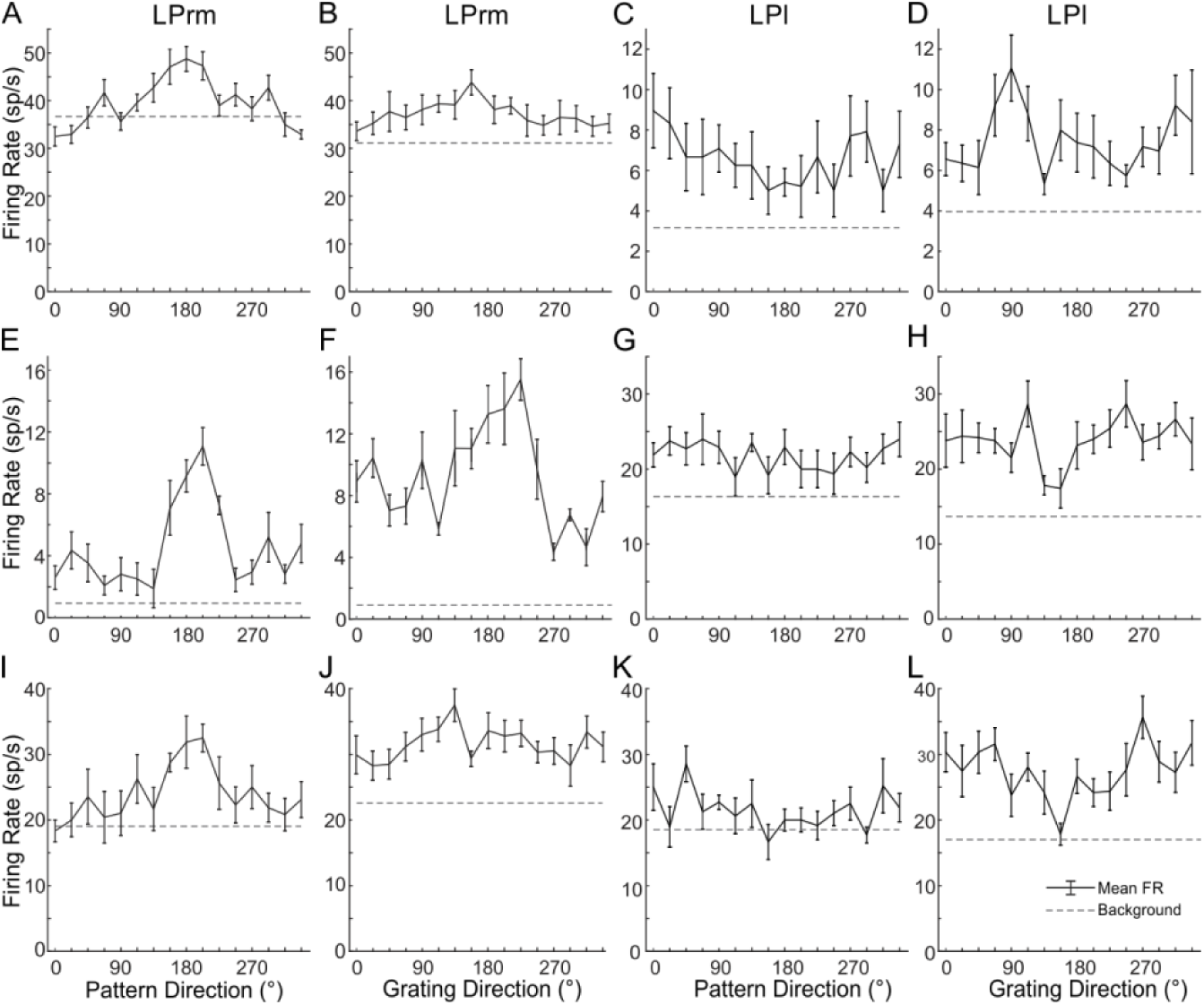
LPrm cells respond to pattern motion, whereas LPl cells do not. Three examples of LPrm cells responding to pattern motion are shown in the first column (A,E,I). The response of these same cells to single gratings is shown in the second column (B,F,J). In contrast, three LPl cells are shown in (C,G,K) which do not respond selectively to pattern motion, but do show selectivity for single moving gratings (D,H,I)

**Figure 8.**
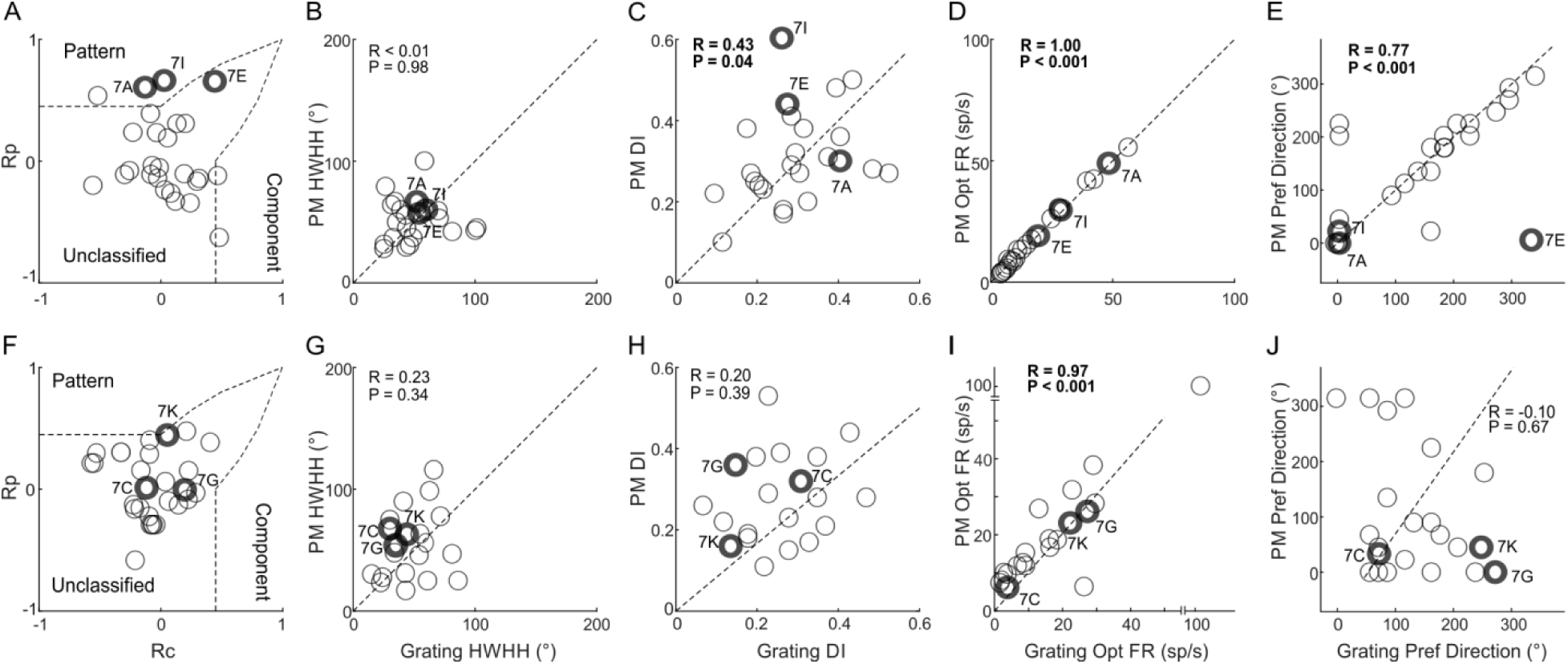
Population summary reveals more pattern motion selectivity in LPrm. Cell distributions for LPrm (top row) and LPl (bottom row) are compared for pattern vs. component selectivity (A,F), pattern motion (PM) vs. single grating HWHH (B,G) and DSI (C,D), peak firing rate (D,I) and preferred direction (E,J).

However, the lack of cells classified as pattern selective may be due to very broad orientation tuning, as was shown in Figure 4AB, which makes comparisons between conditions more difficult. To account for this, we compared quantitative measures of HWHH and direction selectivity in a manner similar to previous studies looking at pattern motion responses in cat LP (Casanova and Savard, 1996). No dependence between HWHH of pattern and single gratings was found for either LP subdivision (Figure 8BG). However, pattern motion sensitivity was detected, specifically in LPrm, using DSI comparisons and the preferred direction itself. The DSI of both pattern motion and single gratings was highly correlated in LPrm (R = 0.43, P < 0.05; Figure 8C), but more weakly correlated and not significant in LPl (R = 0.20, P = 0.40; Figure 8H). Moreover, whereas correlation of preferred direction between pattern and single gratings was high for LPrm (R = 0.77, P < 0.001; Figure 8E), there was no correlation for LPl (R = −0.10, P > 0.60; Figure 8J). In addition, while both LPrm (Figure 8D) and LPl (Figure 8I) showed similar response magnitude to pattern and single grating stimuli, pattern responses were more frequently lower for LPl.

## Discussion

The aim of this project was to test response selectivity in the rat pulvinar/LP. Characteristics of single-unit responses from rostromedial (LPrm) and lateral (LPl) subdivisions of the pulvinar revealed ∼50% of the population responded to drifting gratings, whereas ∼75% was visually responsive. Close to 90% of V1 cells were responsive to both types of stimuli. Further analysis revealed that LPrm and LPl cells had significantly larger receptive field sizes, wider orientation tuning bandwidths, longer response latencies, and higher spontaneous firing rates compared to cells in V1. We also examined differences between pulvinar subdivisions, finding that cells in the colliculo-recipient LPl had a higher mean temporal frequency preference than cells in LPrm. LPrm, on the other hand, showed a greater degree of pattern motion detection. These results suggest that LPl cells may inherit spatiotemporal properties primarily from superior colliculus, whereas LPrm likely derives higher level motion processing through the visual cortex.

### Single Cell Response Characteristics in Rat Pulvinar

The difference between LP and V1 receptive field sizes is indicative of the pulvinar’s role as an integrating higher-order thalamic relay (Guillery and Sherman, 1998). The lateral geniculate nucleus (LGN), a first order relay, receives its driving input directly from the retina, giving its cells receptive fields that are smaller than found in V1 (Rodieck, 1979; Gao et al., 2010). But in pulvinar, receptive fields were large compared to V1; meaning that pulvinar cells may integrate input from many V1 cells, or inherit large receptive field sizes from higher visual cortex, or, in the case of LPl, from the superior colliculus. Relatively large pulvinar receptive fields have also been found in cat, where receptive fields in striate-recipient pulvinar average 8° in diameter (Casanova et al., 1989), versus 2° in V1 (Albus, 1975), and in monkeys, where receptive fields average 5° in diameter in the ventrolateral (Gattass et al., 1979) and dorsomedial pulvinar (Petersen et al., 1985), versus diameters typically less than 1° in V1 (Hubel and Wiesel, 1968). We found V1 receptive field sizes of about 20° on average, similar to previously reported values in rat (Girman et al., 1999; Foik et al., 2018), and LP receptive field sizes averaging 70°. Superior colliculus receptive fields in rat have been reported with diameters ranging from 10° to 60°. Larger receptive field sizes in LP neurons have also been reported in mice (Allen et al., 2016; Bennett et al., 2019; Siegle et al., 2019).

The higher average temporal frequency preference in LPl compared to V1 (Figure 5) suggests some pulvinar neurons might be involved in circuitry to filter visual input based on speed. This would be congruent with the results of Tohmi et al. (Tohmi et al., 2014), who showed that velocity tuning differences between areas in higher visual cortex are lost when superior colliculus is ablated, presumably because pulvinar cells carry the information from temporal-frequency tuned superior colliculus cells which receive input mostly from Y-like and W-like channels (Prévost et al., 2007; Waleszczyk et al., 2007). The longer response latencies we observed in LP compared to V1 (Figure 3) suggest that feedback from spatially tuned visual cortex cells might combine with superior colliculus input in the pulvinar (Jarosiewicz et al., 2012). This conclusion is also supported by previous anatomical studies showing direct inputs from SC to LPl in mouse (Zhou et al., 2017).

Longer latencies observed in our LP cells might also be caused by cortical input alone. Cells in the higher visual cortex of macaque are known to have longer latencies than in V1 on average, although the distribution of latencies within any one particular visual area is large (Schmolesky et al., 1998). In LPrm in particular, which receives no collicular input but does receive cortical projections from V1 and V2 (Takahashi, 1985; Masterson et al., 2009), the additional delay versus V1 likely arises from the addition of synapses from cortex to LP. Similar latency to what we found has been previously reported in V1 of rat (Wang et al., 2006; Foik et al., 2018) and mouse (Durand et al., 2016), but response latencies in some areas of higher visual cortex in mouse are much longer than in V1 on average (Polack and Contreras, 2012), which could account for longer response delays in LP cells.

### Direction selectivity in the LP

Orientation selectivity and direction selectivity have both been demonstrated in rat V1 previously (Girman et al., 1999), and have also been reported in the pulvinar of mouse (Bennett et al., 2019; Siegle et al., 2019), rabbit: (Casanova and Molotchnikoff, 1990; Molotchnikoff and Shumikhina, 1996), cat (Casanova and Savard, 1996; Merabet et al., 1998), and monkey (Petersen et al., 1985; Gattass et al., 2018). Here we confirm that cells in rat LP also respond preferentially to specific orientation and/or direction of drifting sinusoidal gratings, with broader tuning bandwidth compared to V1 cells. This difference between LP and V1 in tuning width is consistent with studies performed in cats (Casanova et al., 1989; Chalupa and Abramson, 1989). We also found that, unlike V1, LP cells can respond selectively to pattern motion stimuli as has been shown previously in pulvinar of cats (Merabet et al., 1998, 2000) and monkeys (Chalupa et al., 1976; Benevento and Miller, 1981). Component-pattern motion analyses previously applied to primate higher cortical visual areas (Movshon et al., 1985; Movshon and Newsome, 1996) revealed only a few pattern selective cells in LPrm. However, such analysis was made difficult here due to the nearly twice as broad tuning widths of LP cells compared to neurons in visual cortex. When making comparisons using analyses previously done on cat LP-pulvinar complex neurons (Casanova and Savard, 1996) we found a strong correlation of direction response between pattern motion and single drifting gratings, again suggesting its potential role in processing motion. This result was specific to LPrm and can be used to distinguish it from LPl. This also suggests a stronger input from higher visual cortex to LPrm, as this is where pattern motion has been shown to emerge in rodent (Juavinett et al., 2015) and other species (Movshon et al., 1985; Casanova and Savard, 1996; Movshon and Newsome, 1996).

In summary, our findings suggest that rat pulvinar is comparable to the pulvinar of other mammals in terms of visual processing. We verified the presence of orientation and direction tuned cells in LP, and identified receptive fields with large diameters compared to V1 cells. We also distinguished two distinct roles for the rostromedial and lateral portions of LP. The lateral part is more likely involved in temporal frequency and speed processing, whereas the rostromedial portion has stronger detection of pattern, perhaps due to differences in collicular input or connectivity with higher visual cortex. Further research aiming to identify differences in cortical connectivity with pulvinar subdivisions will help answer outstanding questions regarding the anatomical routes by which pulvinar might derive spatiotemporal receptive field properties.

